# Topographical features enhance DNA tetrahedron biostability

**DOI:** 10.64898/2026.10.27.762189

**Authors:** Sarah Youssef, Lauren A. Anderson, Mengyao Zheng, Arun Richard Chandrasekaran

## Abstract

Stabilization of nucleic acid-based systems is critical for their clinical translation in biomedical applications. Here, we integrated hairpins within the edges of a DNA tetrahedron and studied the stabilization induced by these motifs against nucleases. We show that nuclease resistance of DNA tetrahedra increased with the number of hairpins in the structure, this biostability enhancement is specific to the sequences within the hairpin. Our study provides a new design approach to create nucleic acid nanostructures with inherent biostability.

---

Being a natural material, DNA nanostructures have shown the most promise in biomedical applications such as diagnostics, bioimaging and drug delivery. One of the key impediments in realizing this potential is the insufficient stability of DNA nanostructures in biological environments.^1–3^ Several strategies have been developed to enhance the biostability of DNA nanostructures including the incorporation of chemical modifications,^4^ polymer coatings^5^ and crosslinking of DNA strands.^6^ Design-based biostability of DNA nanostructures^7^ and the role of size^8^ and shape have also been investigated. Enhancing the nuclease resistance of DNA nanostructures provides additional characteristics for these structures to be better suited for biological applications.

In this study, we studied the stabilizing effect of hairpins with different sequences incorporated within the edges of a DNA tetrahedron. DNA tetrahedra are one of the widely used DNA nanostructures in biological applications, with the geometry of the structure allowing efficient cellular internalization.^9,10^ Our study stems from prior literature that showed high nuclease resistance of specific hairpin sequences.^11,12^ The enhanced nuclease resistance of the hairpin sequence GCGAAGC was attributed to the stacking effect created by additional hydrogen bond formation between two adenine bases and guanine base. This phenomenon prevented nucleases, particularly exonucleases from accessing the phosphodiester bond at the end of DNA strands. We anticipated that incorporating such hairpins within DNA tetrahedra would increase their resistance against nucleases. We evaluated the effects of hairpin sequence, number and spatial organization on the overall stability of tetrahedra against nucleases.

We inserted hairpins along the edges of the tetrahedron and varied the number of hairpins and the sequence. The model DNA tetrahedron we used contains 20 bp per edge and is composed of four strands.^13^ To modulate the number of hairpins in the structure, we modified a subset of strands to contain a single hairpin, enabling us to construct DNA tetrahedra with 1, 2, 3 or 4 hairpins (**Fig. 1a**). The first hairpin we used was a GCGAAGC sequence with a GC:GC stem and a GAA loop. To evaluate the effect of the hairpin sequence, we designed two other insertions on the tetrahedra edges: (1) a hairpin sequence with the same GC:GC stem but with TTT in the loop since a GC:GC stem is known to be more stable than an AT:AT stem^14^ and (2) a poly T sequence that will be a single stranded loop instead of a hairpin (**Fig. 1b**). The hairpins were designed to be located in the middle of the edge in the DNA tetrahedron, with two of the hairpins placed at the nick already present on the tetrahedral edge and the other two inserted within a strand. We first confirmed proper assembly of the structures using nondenaturing polyacrylamide gel electrophoresis and observed results similar to prior studies on DNA tetrahedra assembly.^8,15^ The structures with different number of hairpins were formed with minimal loss in assembly yields compared to the structure without any hairpins (**Fig. 1b** and **Fig. S1**).

**Figure 1.**
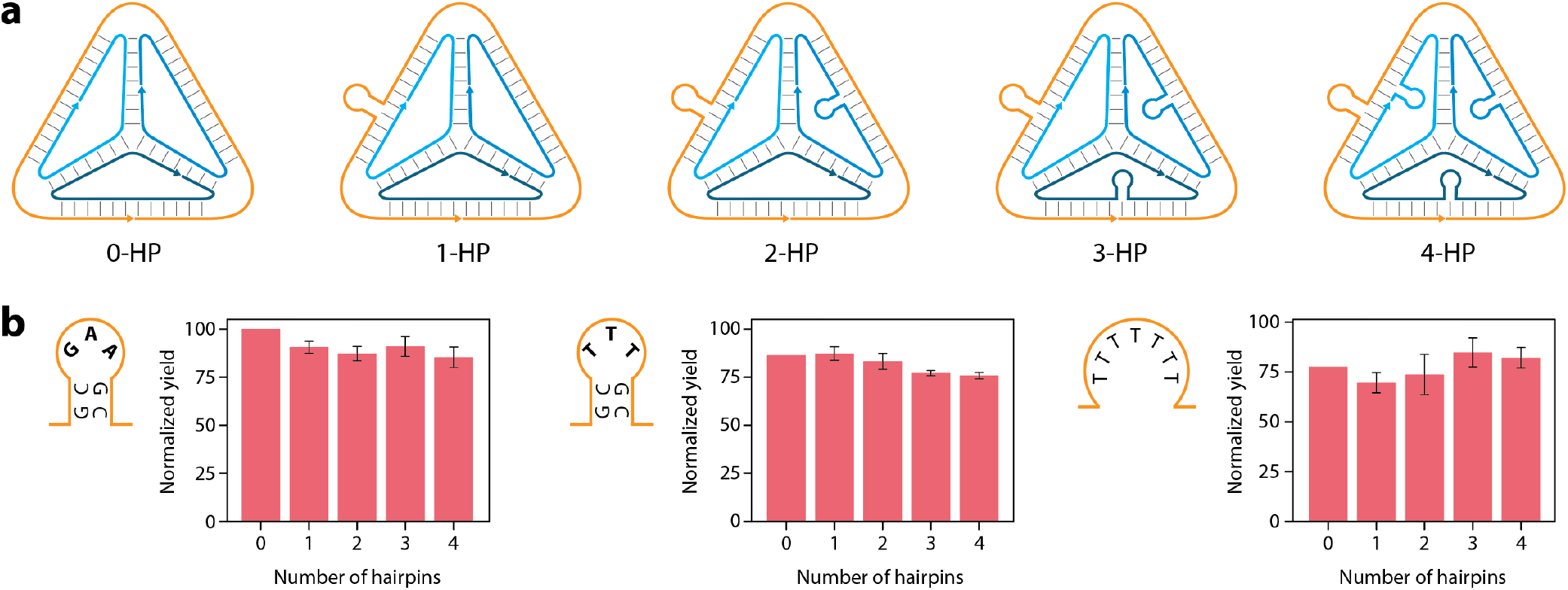
(a) Scheme of DNA tetrahedra with different number of hairpins. The tetrahedron contains four strands and one hairpin is incorporated into each strand to obtain structures with more hairpins. (b) Assembly yields of tetrahedra with different number of hairpins for each hairpin sequence: a GAA sequence in the loop, a TTT sequence in the loop and a fully polyT sequence instead of a hairpin.

To test nuclease resistance, we first chose DNase I, one of the most abundant nucleases in human serum.^16^ We tested different amounts of DNase I for the GCGAAGC hairpin sequence with different number of hairpins on the tetrahedron. To eliminate any effect of the additional base pairs on DNase I activity, we normalized the concentration of the DNA tetrahedra with different number of hairpins used to study DNase I activity. We observed that the nuclease resistance increased with the number of hairpins, with the 4-hairpin structure showing the highest nuclease resistance (**Fig. 2a** and **Fig. S2**). The tetrahedron with four hairpins was ∼60% intact against 0.3 U DNase I while tetrahedra with lower number of hairpins were less than ∼30% intact at the same enzyme amount.

**Figure 2.**
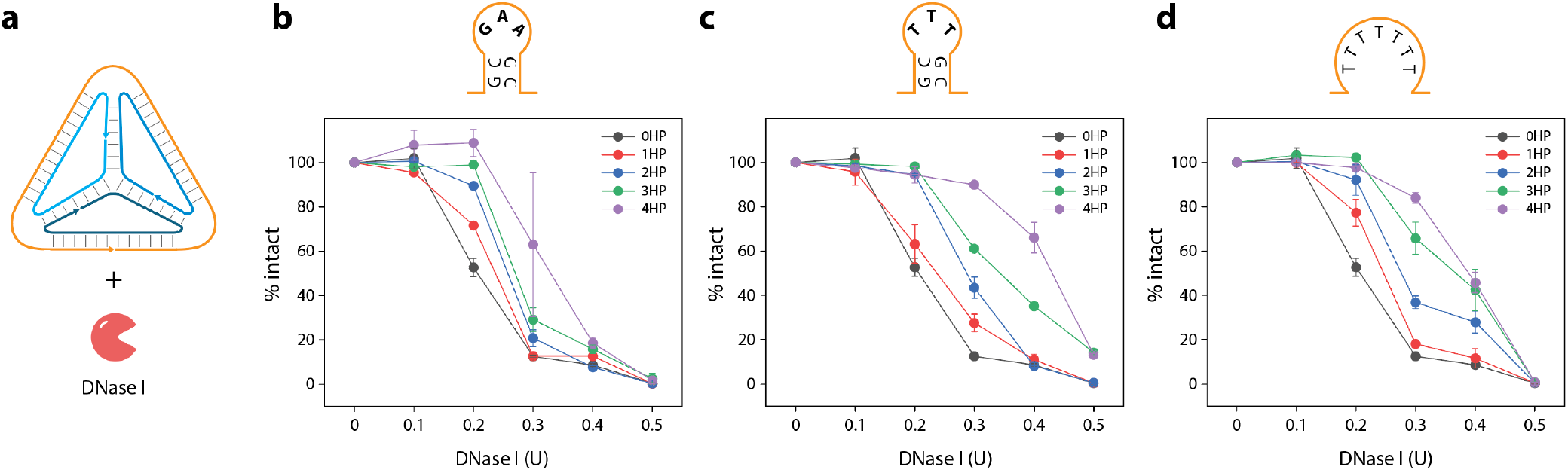
(a) Nuclease resistance analysis against DNase I. (b-d) Degradation trends of tetrahedron with GAA loop, TTT loop and a poly T sequence, respectively. Data and error bars shown are average and standard deviation values from replicate experiments.

Next, we studied the distribution of hairpins among the four strands that form the tetrahedron. Instead of each strand containing a single hairpin, we tested tetrahedra where only one of the component strands was modified to contain 1, 2, 4 or 6 hairpins (**Fig. S3**). While the assembly yields slightly varied, the structure was overall well-formed when containing up to 4 hairpins on one of the strands. The structure containing 6 hairpins did not assemble well and was omitted from the rest of the study (**Fig. S3**). This could be due to the increasing number of well-stacked hairpins obstructing the kinetically favored hybridization reaction during the assembly of the tetrahedron containing 6 hairpins. When tested against DNase I, we again observed that the 4-hairpin structure was the most nuclease resistant (**Fig. S4**). However, the assembly yields of the DNA tetrahedron when multiple hairpins were inserted into the same strand was inconsistent compared to that of the tetrahedron where each of the 4 strands contained a single hairpin each (totaling 4/structure). This result suggests that distributing hairpins around strands to cover larger area of tetrahedron was more effective in both the assembly as well as in shielding the structure against DNase I. Thus, we chose the structure with 4 hairpins with one hairpin per strand for further experiments (**Fig. S3**).

Next, we tested whether the sequence of the hairpin head was crucial for conferring nuclease resistance using the TTT head sequence and a T_7_ loop instead of a hairpin (**Fig. 2b-c** and **Fig. S2**). We observed that the hairpin sequence with the TTT head showed higher nuclease resistance compared to the hairpin with the GAA head or the T_7_ loop. This was in contrast to previous reports that the hairpin sequence containing a GAA head was highly biostable.^11,12^ Poly-T stretches are commonly used in DNA origami structures to circumvent end helical stacking since they create an electron cloud forming a repulsion zone.^17^ We assumed a similar effect between the TTT head and DNase I leads to higher degree of stabilization, a phenomenon that requires further investigation in the future.

Since the hairpins were originally studied against exonucleases, we tested if incorporation of such hairpins within tetrahedra will stabilize them against exonucleases. We ran experiments against commonly abundant exonucleases T5 Exo and Exo III. In contrast to DNase I, T5 exonuclease did not show any enhancement in nuclease resistance with the hairpins (**Fig. 3a** and **Fig. S5**). When tested against exonuclease III, the tetrahedra with four polyT loops and TTT head showed higher nuclease resistance compared to the structure without any hairpins (**Fig. 3b** and **Fig. S5**). The hairpin with the GAA head hairpin only provided moderate enhancement.

**Figure 3.**
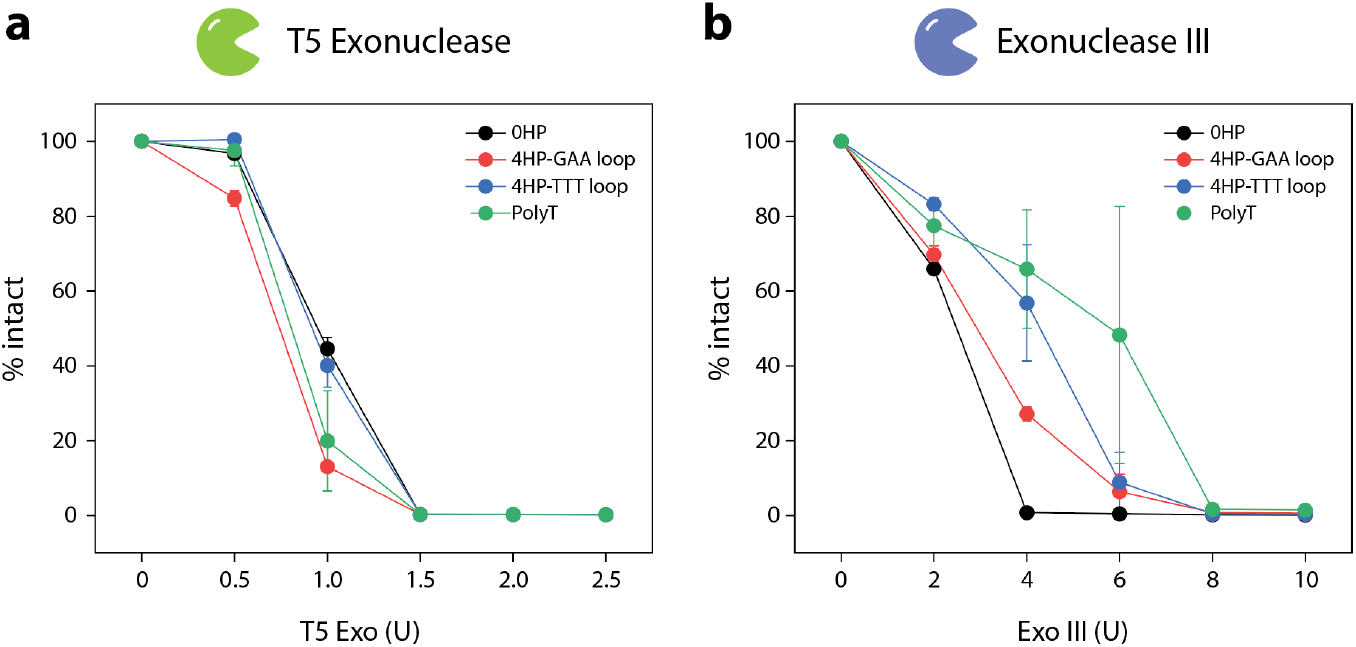
Degradation trends of tetrahedron with four hairpins compared against tetrahedra without any hairpins in (a) T5 exonuclease and (b) Exonuclease III. Data and error bars shown are average and standard deviation values from replicate experiments.

Overall, our results show that strategic incorporation of short hairpins within the edges of DNA tetrahedra enhances the nuclease resistance of the structure. Our study informs on the distribution of the hairpins on the structure as well as the choice of hairpin sequences for enhanced biostability. However, a dominant effect of the type of sequence within the hairpin across different nucleases is still uncertain, and a more systematic analysis of the stem and loop sequences is required. This strategy can also be implemented in different sizes of DNA tetrahedra to ascertain whether the loss in biostability due to size^8^ can be compensated by the inclusion of hairpins on the edges. Our work adds to the knowledge base for biostability enhancement of DNA nanostructures, leading to their use in biomedical applications.

## Competing interests

The authors have no competing interests.

## Supporting information

Experimental procedures, additional results and DNA sequences used.

## Author contributions

S.Y. conceived the project, designed experiments, analyzed data and wrote the paper. L.A. and M.Z. performed experiments and analyzed data. A.R.C. supervised the project, designed experiments, analyzed data, visualized data and wrote the paper.

## Acknowledgments

Research reported in this publication was supported by the National Institutes of Health (NIH) through National Institute of General Medical Sciences (NIGMS) under award number R35GM150672 to A.R.C.

## Notes

### Competing Interest Statement

The authors have declared no competing interest.

