## Supporting information for "Topographical features enhance DNA tetrahedron biostability"

### MATERIALS AND METHODS

#### Preparation of DNA complexes

DNA strands were purchased from Integrated DNA Technologies (IDT). Full sequences are listed in Table S1. DNA tetrahedra were prepared by mixing the component DNA strands in equal ratios in Tris-Acetic-EDTA buffer containing 40 mM Tris base (pH 8), 20 mM acetic acid, 2 mM EDTA, and 12.5 mM magnesium acetate (1× TAE-$Mg^{2+}$). Samples were annealed by incubating at 90 °C for 5 minutes using a heating block, followed immediately by incubation on ice for 5 minutes.

#### Nuclease degradation assay

DNA tetrahedra were first mixed with the appropriate reaction buffers provided with the enzymes (final of 1×). The concentration of the DNA tetrahedra with differing numbers of hairpins was normalized to the number of base pairs in the structure. Enzyme dilutions were made in nuclease-free water. For the nuclease degradation assay, 1 µl of the enzyme was added to 9 µl of the sample containing DNA tetrahedron and reaction buffer. Samples were incubated at 37 °C for 30 min.

#### Gel electrophoresis and quantification

Non-denaturing gels were prepared using 19:1 acrylamide/bisacrylamide (National Diagnostics). Samples were mixed with loading dye containing bromophenol blue and glycerol prior to loading. Gels were run at a constant voltage at 4 °C in 1× TAE-$Mg^{2+}$ running buffer. Gels were stained in 0.5× GelRed (Biotium), imaged using a Bio-Rad Gel Doc XR+ and analyzed using ImageLab software. Images were typically taken at multiple exposures to facilitate accurate quantification. For each gel, quantification was done using the highest-exposure image that did not contain saturated pixels in the bands. For analysis of replicates, separate gels with the same exposure time were typically used for quantifying band brightness. The assembly yield was quantified as the relative intensity of the tetrahedron band with hairpins to that of the tetrahedron without any hairpins. Nuclease degradation profiles were obtained by normalizing the band intensity corresponding to the structure (at each enzyme concentration) to the control lane without any enzyme.

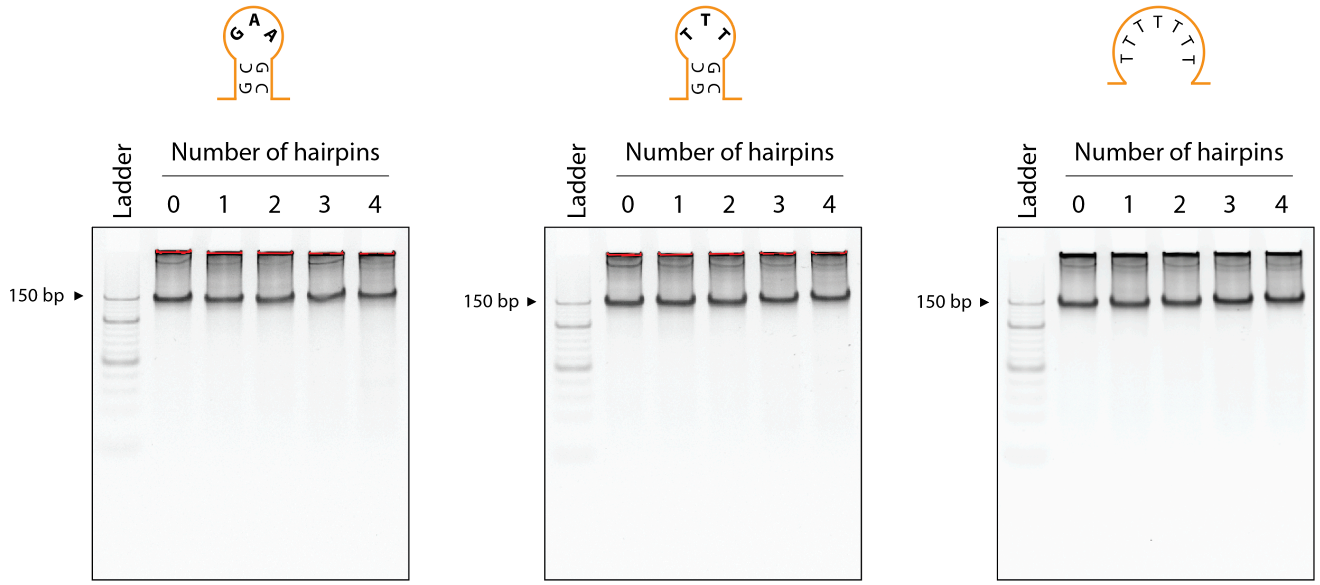


**Figure S1**. Non-denaturing gels showing assembly of DNA tetrahedra.

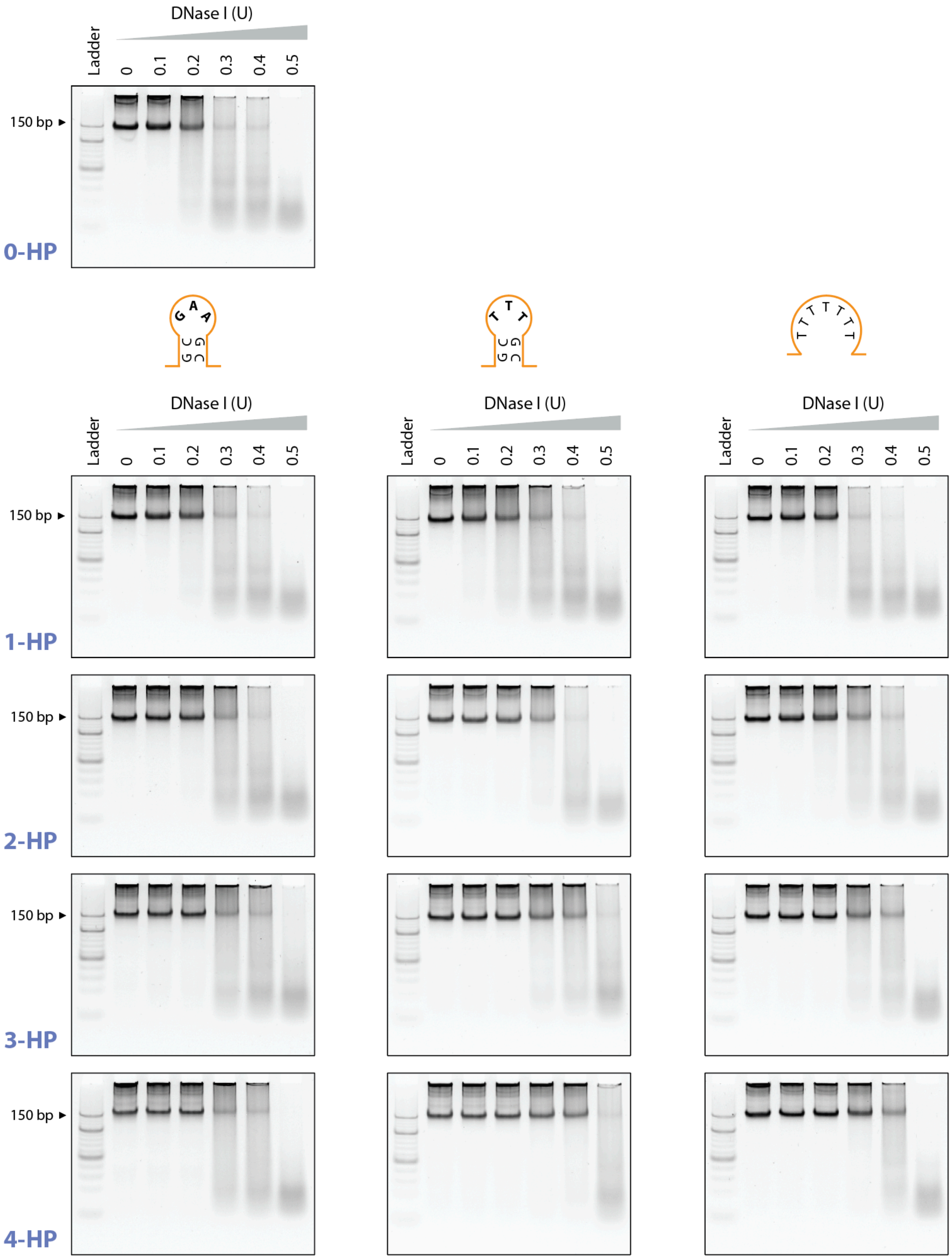


**Figure S2**. Non-denaturing gels showing degradation of DNA tetrahedra when incubated with DNase I.

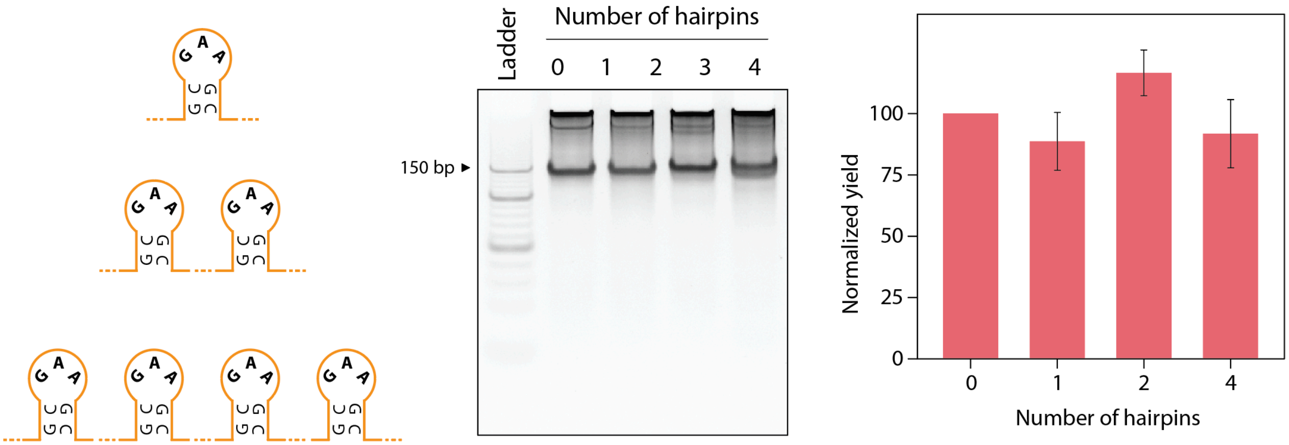


**Figure S3**. Non-denaturing gels showing assembly of DNA tetrahedra with multiple hairpins on one strand.

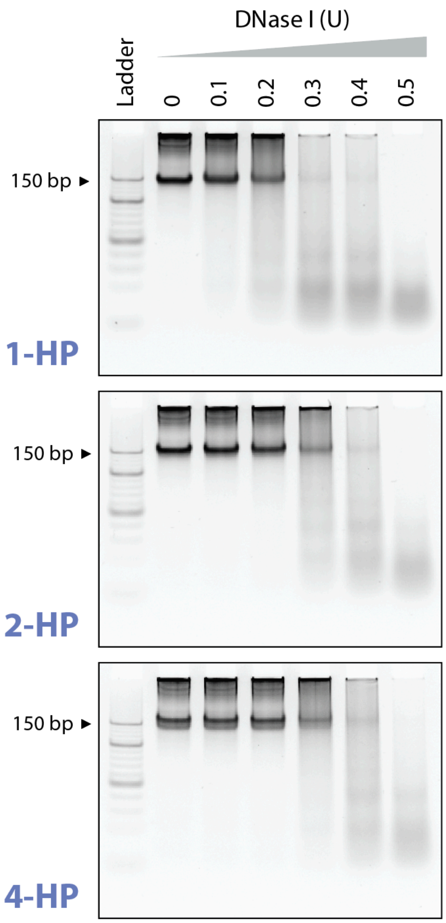


**Figure S4**. Non-denaturing gels showing degradation of DNA tetrahedra with multiple hairpins on one strand when incubated with DNase I.

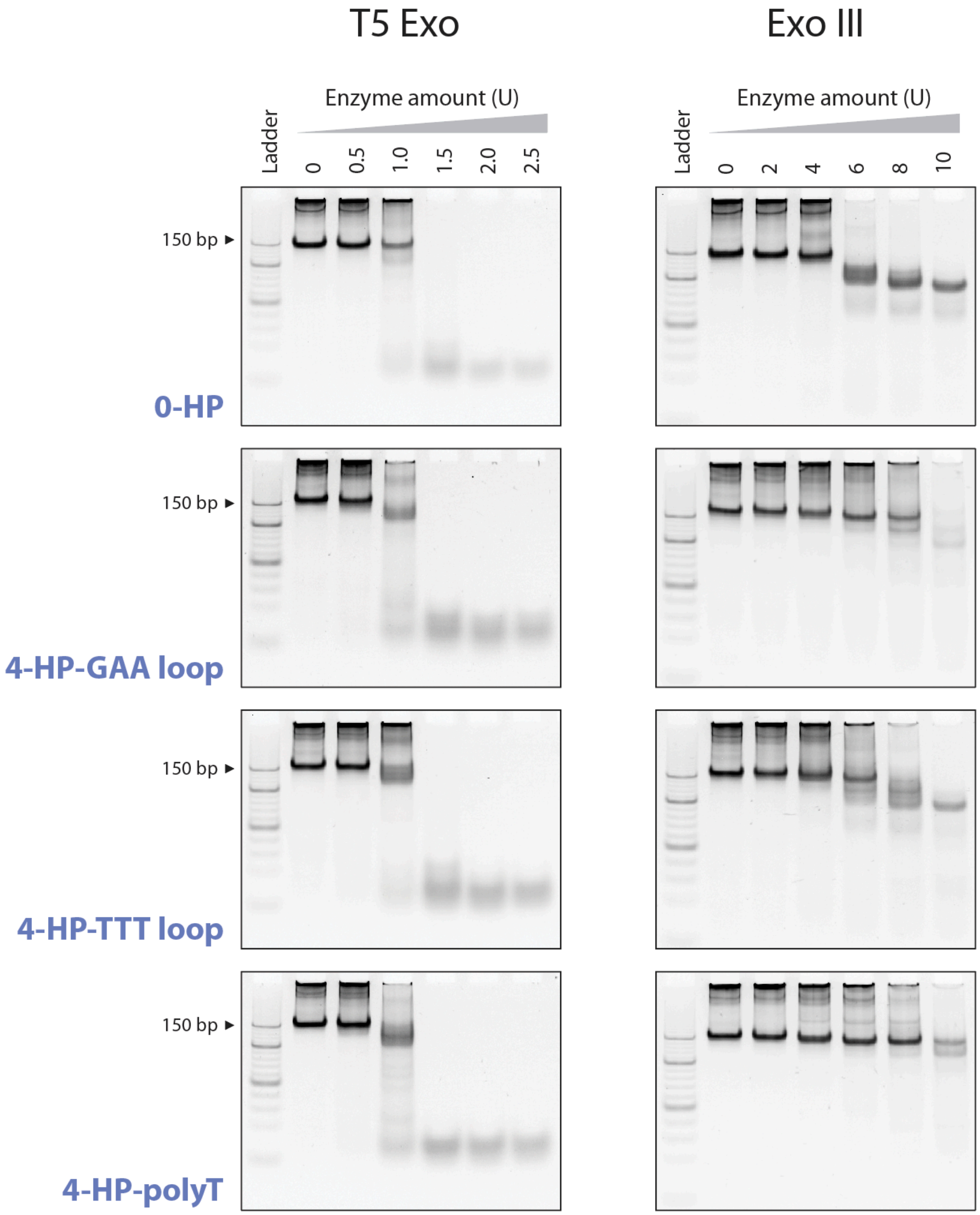


**Figure S5**. Non-denaturing gels showing degradation of DNA tetrahedra when incubated with exonucleases.

| Strand name | Sequence (5’ to 3’) |
| --- | --- |
| S1 | AGGCAGTTGAGACGAACATTCCTAAGTCTGAAATTTATCACCCGCCATAGTAGACGTATCACC |
| S2 | CTTGCTACACGATTCAGACTTAGGAATGTTCGACATGCGAGGGTCCAATACCGACGATTACAG |
| S3 | GGTGATAAAACGTGTAGCAAGCTGTAATCGACGGGAAGAGCATGCCCATCCACTACTATGGCG |
| S4 | CCTCGCATGACTCAACTGCCTGGTGATACGAGGATGGGCATGCTCTTCCCGACGGTATTGGAC |
| S1-1HP-GAA loop | AGGCAGTTGAGACGAACATTCGCGAAGCCTAAGTCTGAAATTTATCACCCGCCATAGTAGACGTATCACC |
| S2-1HP-GAA loop | CTTGCTACACGATTCAGACTTAGGAATGTTCGACATGCGAGGGTCCAATACCGACGATTACAGGCGAAGC |
| S3-1HP-GAA loop | GGTGATAAAACGTGTAGCAAGCTGTAATCGACGGGAAGAGCATGCGAAGCGCCCATCCACTACTATGGCG |
| S4-1HP-GAA loop | CCTCGCATGACTCAACTGCCTGGTGATACGAGGATGGGCATGCTCTTCCCGACGGTATTGGACGCGAAGC |
| S1-1HP-TTT loop | AGGCAGTTGAGACGAACATTCGCTTTGCCTAAGTCTGAAATTTATCACCCGCCATAGTAGACGTATCACC |
| S2-1HP-TTT loop | CTTGCTACACGATTCAGACTTAGGAATGTTCGACATGCGAGGGTCCAATACCGACGATTACAGGCTTTGC |
| S3-1HP-TTT loop | GGTGATAAAACGTGTAGCAAGCTGTAATCGACGGGAAGAGCATGCTTTGCGCCCATCCACTACTATGGCG |
| S4-1HP-TTT loop | CCTCGCATGACTCAACTGCCTGGTGATACGAGGATGGGCATGCTCTTCCCGACGGTATTGGACGCTTTGC |
| S1-1HP-polyT | AGGCAGTTGAGACGAACATTCTTTTTTTCTAAGTCTGAAATTTATCACCCGCCATAGTAGACGTATCACC |
| S2-1HP-polyT | CTTGCTACACGATTCAGACTTAGGAATGTTCGACATGCGAGGGTCCAATACCGACGATTACAGTTTTTTT |
| S3-1HP-polyT | GGTGATAAAACGTGTAGCAAGCTGTAATCGACGGGAAGAGCATTTTTTTTGCCCATCCACTACTATGGCG |
| S4-1HP-polyT | CCTCGCATGACTCAACTGCCTGGTGATACGAGGATGGGCATGCTCTTCCCGACGGTATTGGACTTTTTTT |
| S1-2HP-GAA loop | AGGCAGTTGAGGCGAAGCACGAACATTCCTAAGTCTGAAGCGAAGCATTTATCACCCGCCATAGTAGACG TATCACC |
| S1-4HP-GAA loop | GCGAAGCAGGCAGTTGAGGCGAAGCACGAACATTCCTAAGTCTGAACGGAAGCATTTATCACCCGCCATA GTAGGCGAAGCACGTATCACC |
| S1-6HP-GAA loop | GCGAAGCAGGCAGTTGAGGCGAAGCACGAACATTCGCGAAGCCTAAGTCTGAAGCGAAGCATTTATCACC GCGAAGCCGCCATAGTAGGCGAAGCACGTA |

**Table S1.** Sequences used in the study (written 5' to 3').
